# A default-control network cortical gradient differentiates the imagination of social and solitary experiences

**DOI:** 10.1101/2025.11.25.690561

**Authors:** Andrew J. Anderson, Adam G. Turnbull, Jonathan Smallwood, Feng V. Lin

**Affiliations:** Neurology, Medical College of Wisconsin, WI, USA; Biomedical Engineering, Medical College of Wisconsin, WI, USA; Neurosurgery, Medical College of Wisconsin, WI, USA; Wu Tsai Neurosciences Institute, Stanford University, USA; Department of Psychology, Queens University, CA; Department of Psychiatry and Behavioral Sciences, Stanford University, USA

## Abstract

Understanding the neural basis of spontaneous thought - when attention shifts from external tasks to internally generated content such as reminiscing or planning - remains a central challenge in cognitive neuroscience. Progress in this area has lagged studies of externally driven cognition, in part because self-generated thought is difficult to control and measure. Recent work suggests that transitions between externally and internally focused cognitive states follow a continuous neural activation gradient that reflects opposing patterns of engagement of the frontoparietal control network versus the default mode network. To characterize internally focused cognitive states that prospectively engage the default mode network, we used functional Magnetic Resonance Imaging (fMRI) to measure brain activity while participants imagined a range of personal scenarios prompted by generic text cues (e.g., party, housework) to mimic natural thought. Before scanning, participants described each scenario verbally and rated its experiential feature content. Gradient-space analysis revealed a cognitive transition within imagined states: solitary activities (e.g., housework) preferentially recruited the frontoparietal network relative to the default mode network, whereas social activities (e.g., a party) showed the opposite pattern. Because all cognitive states were internally focused (imagined) and elicited via the same non-interactive task, these results refine interpretations of the frontoparietal-default mode gradient. They show that this gradient does not simply differentiate external task-positive from internal task-negative states but also tracks semantic differences between internal cognitive states.

## Introduction

Functional Magnetic Resonance Imaging (fMRI) has revealed important insights about how large-scale brain networks support different cognitive states. Although most studies focus on carefully designed, interactive external tasks, fMRI has also been used to examine internally generated cognitive states associated with mind wandering and imagination (Smallwood & Schooler, 2015; Smallwood et al., 2021). These spontaneous thoughts occupy an estimated 30–50% of waking life (Killingsworth & Gilbert, 2010) and are central to both cognition (Yanko & Spalek, 2014; Smallwood et al., 2007) and emotional well-being (Marchetti et al., 2016). However, they remain difficult to study with neuroimaging because internal states are self-generated, difficult to control, and require self-reporting, which can disrupt ongoing thought, and be unreliable (Kane et al., 2021). Understanding the neural basis of these internal states is therefore a central challenge in cognitive neuroscience.

Early work showed that cognitive states can be distinguished by regional and network-level activation patterns (Fox et al., 2005) and internally focused processes such as reminiscing, conceptual thinking, and planning primarily engaged the default mode network (DMN; Binder et al., 1999; Mazoyer et al., 2001), whereas externally focused, goal-directed interactive tasks recruited the frontoparietal control network (FPN; Duncan, 2010; Fedorenko et al., 2013). This distinction led to the influential “task-negative” versus “task-positive” framing of DMN and FPN activation/deactivation (Fox et al., 2005), with the degree of (dis)connectedness indicative of task performance (e.g. Hampson et al. 2010). Later studies, however, revealed that these networks can cooperate as well as compete (Christoff et al. 2009, Spreng et al., 2012). Internally generated but goal-oriented thinking, for example, increases co-activation and connectivity between the DMN and FPN (Spreng et al., 2010; Smallwood et al., 2012), suggesting FPN supports goal structure, regardless of whether the goal concerns internal or external content, whereas DMN supports internally focused thought.

More recently, researchers have emphasized whole-brain activation patterns that reflect cooperation and competition between large-scale networks, characterized using functional gradients derived from factor analyses of resting-state data (Margulies et al., 2016; Huntenburg et al., 2018; Smallwood et al., 2021). These gradients capture smooth transitions in activation across the cortex, and one prominent gradient spans a continuum from DMN to FPN related states (Margulies et al., 2016; Karapanagiotidis et al., 2020; McKeown et al., 2020). During tasks, brain state transitions along this gradient axis account for substantial variance in brain activity and relate to cognitive abilities such as fluid intelligence (Shine et al., 2019). They also track shifts between externally focused task engagement and internally oriented off-task states measured with thought probes (Turnbull et al., 2020) and align with distinctions between vigilance and target detection (McKeown et al., 2023).

Despite these advances, relatively little is known about how the semantic representation of internal cognitive states, and not just their internal/external orientation, are expressed in whole-brain gradient space. Approaches using thought probes (Christoff et al., 2009; Sormaz et al., 2018), trait measures (Gorgolewski et al., 2014; Smallwood & Schooler, 2015), or structured internally-oriented tasks (Andrews-Hanna et al., 2010, 2014; Zhang et al., 2022; Shao et al., 2024) provide valuable insights, they do not directly map the semantic structure of thought to these large-scale brain patterns. Continuous verbal reporting (Gilmore et al., 2021; Li et al., 2023; Su et al., 2025) offers richer access to mental content but may disrupt natural cognition, and spontaneous thoughts may unfold at timescales that speech cannot capture.

In contrast, the semantic representations evoked by language are relatively well characterized by semantic language models that predict neural responses to linguistic stimuli (Mitchell et al., 2008; Wehbe et al., 2014; Huth et al., 2016; Anderson et al., 2017; Tang et al., 2023). Because linguistic input drives neural activity, these methods can map brain responses onto semantic model vector spaces capturing word meaning that are time-aligned to word stimuli (Pennington et al., 2014; Binder et al., 2016; Radford et al., 2019).

Here, we extend semantic modeling to understand internally generated imagination in gradient space (**Fig. 1**). We use a design approximating naturally occurring internally focused thought while systematically varying semantic content (Anderson et al., 2020). Participants first imagined personal experiences from their memory of twenty scenario cues spanning diverse experiential content (ranging from Party to Housework). They rated each scenario on sensory, motor, social, affective, and spatiotemporal features and provided verbal descriptions. Then, during fMRI, they re-imagined these individualized scenarios. By projecting fMRI data onto canonical resting-state gradients (Hardikar et al., 2024) and mapping gradient-space activity onto semantic vector spaces, we linked the DMN–FPN gradient to cognitive state transitions. Imagining solitary activities, focused on non-human objects, and including goal-directed tasks such as housework, activated FPN relative to DMN, whereas imagining social activities, showed the opposite pattern.

**Figure 1.**
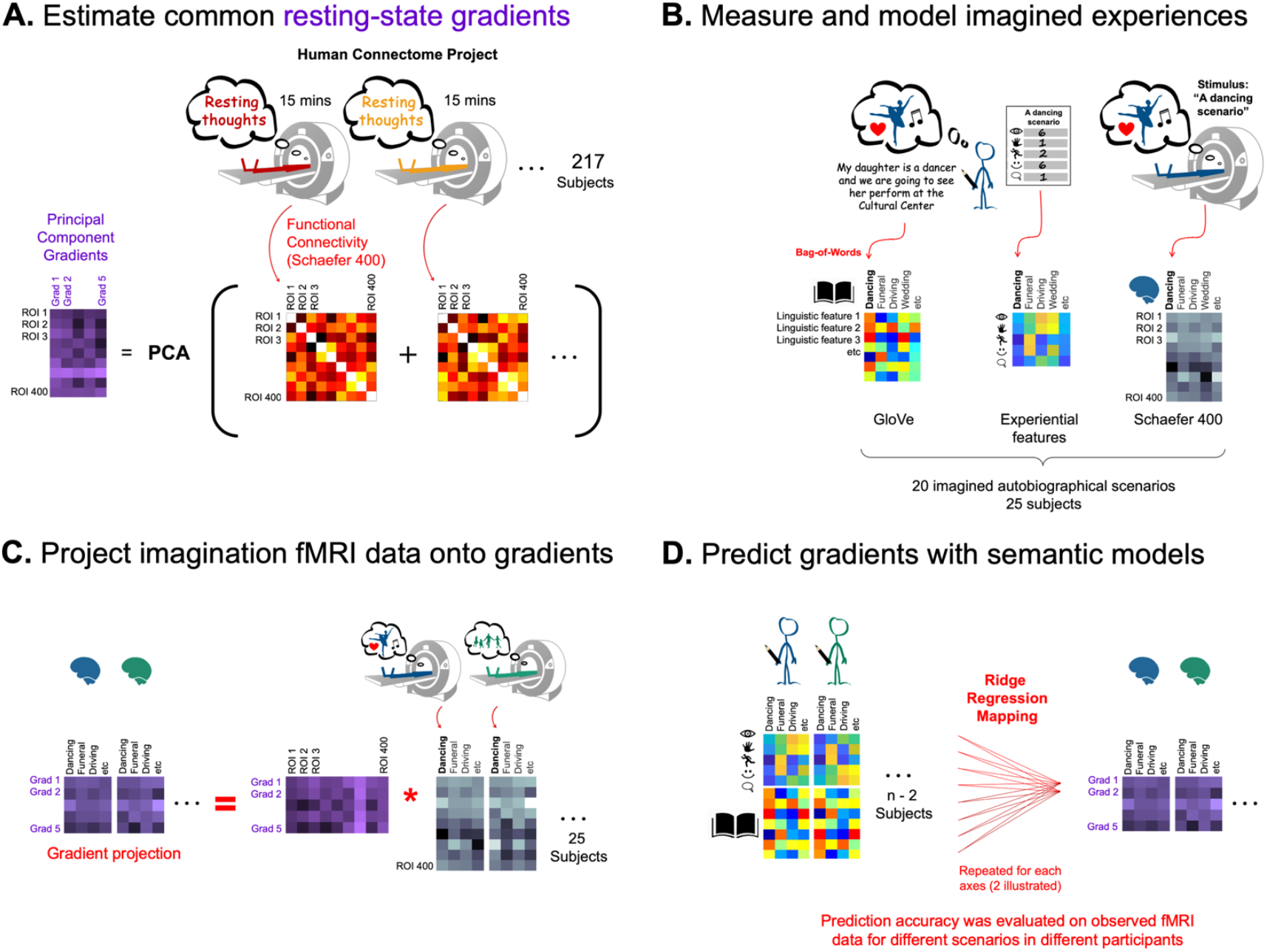
Predicting semantic variation along gradient projections of imagination fMRI data. **A.** Brain activity gradients were estimated by a Principal Components Analysis decomposition of group-average resting-state functional connectivity matrices, represented in Schaefer-400 space. **B**. A separate group of 25 participants imagined themselves personally experiencing 20 scenarios (e.g. “A dancing scenario”), actively simulating their perception, action and feelings. Participants then verbally described each of their imagined scenarios and further rated them [0-6] in terms of twenty sensory, motor, spatial, social, affective and spatiotemporal features of experience. Linguistic descriptions were numerically represented by mapping content words to GloVe word embeddings and then summing word vectors. This was repeated for each scenario and participant. Participants then underwent fMRI as they reimagined the same 20 scenarios, one by one, when cued by written prompts. **C**. To establish whether gradients were systematically modulated during self-generated mental imagery, the corresponding fMRI data was normalized into Schaefer-400 space and projected onto the first five cortical gradients (from **A**). **D** To link gradient activation to participants’ verbal/rating-based descriptions of their imaginary scenarios, we undertook a cross-validated multiple regression analysis. Cross-participant encoding mappings from models to each of the five gradients were fit using ridge regression. Mappings were fit on a concatenation of n-2 participants fMRI data corresponding to 50% of the scenarios. The regularization penalty was selected based on predicting participant n-1’s fMRI data with their personal models for the same 50% of scenarios. The selected encoding mapping was then evaluated on predicting the other 50% of scenarios in the final held-out participant. This analysis was repeated 25 times treating each participant as the test participant. The entire analysis was repeated 100 times using different randomized 50% train vs 50% test scenario splits. Prediction accuracies were averaged across the repeats to produce a final metric for each participant for each gradient, as displayed in **Fig. 2**.

Critically, these findings refine how the DMN–FPN gradient should be interpreted. Because all imagined scenarios were elicited by the same non-interactive task, the gradient cannot simply reflect a task-positive to task-negative axis. Nor can it be reduced to an external–internal distinction, as cognitive states were internally generated by imagination. Instead, the gradient also tracks systematic differences in the semantic content of internal imagined states, cutting across the conventional task-positive/task-negative divide. This reframing sharpens theoretical accounts of functional gradients and clarifies their role in shaping the structure of imaginative thought.

## Results

To test the overarching hypothesis that modulation of brain activity in resting-state functional gradient space (**Fig. 1A**) reflects variation in the semantic content of imagined personal experience, we analyzed a dataset (Wang et al. 2024) comprising twenty-five young adults (Mean±SD age=24±3y, 16F, 9M) who underwent the fMRI experiment illustrated in **Fig. 1B**. Participants undertook a standardized task, where, before scanning, they imagined their idiosyncratic experiences of 20 fixed experimenter-selected scenarios. The scenarios were common, and diverse, spanning a range of events that one plausibly might think about at rest, including social activities such as a party, special occasions such as a wedding, tasks such as housework, and more passive activities such as reading (Anderson et al. 2020).

To enable computational semantic modeling of the idiosyncratic scenarios, participants verbally described each imagined scenario (Anderson et al. 2020) and rated the importance of twenty sensory, motor, affective, social, cognitive and spatiotemporal features of experience to their mental image on a scale of [0 6]. Details of the ratings are contained in **Supplementary Table 1. Supplementary Figs. 1** and **2** provide supporting visualizations of participants’ experiential ratings and word clouds of scenario descriptions. To further characterize the nature of the imagined scenarios and gauge the extent to which they reflected episodic memories (Tulving et al. 1972, Wheeler et al. 1997), participants rated each scenario on whether it reflected a real-life event and whether it was vividly imagined. Ratings indicated that imagined scenarios largely reflected real events (overall mean across scenarios and participants=5.6, scale [0 6]) that were vividly imagined (overall mean=4.6, scale [0 6]).

Participants then underwent fMRI as they re-imagined the same idiosyncratic scenarios, five times over, in random order, when prompted by a generic text prompt (e.g. “A party scenario”). fMRI data were preprocessed with standard methods, including fMRIPrep (Esteban et al. 2019) to produce a single fMRI representation associated with each scenario per participant.

To estimate whether functional gradients reflected systematic changes in the semantic content of scenarios, the imagination fMRI time series were projected onto the first five principal gradients with a matrix multiplication. This yielded five time-series, one per gradient. The critical DMN-FPN gradient was gradient #3, and the other four gradients were included to explore whether they also reflected semantic variation.

The gradients themselves were principal components derived using the BrainSpace toolbox (Vos de Wael et al. 2020) from precomputed group-average functional connectivity matrices of Human Connectome Project resting-state data (van Essen et al. 2013). The resting-state data were from 217 healthy young adults (Mean±SD age=28.5±3.7y, 122F, 95M), preselected in Vos de Wael (2018). Vos de Wael 2018 derived group-level functional connectivity matrices by: (1) Representing the resting-state fMRI data according to the Schaefer-400 parcellation scheme (Schaefer et al. 2018), and pointwise averaging activation across participants, to generate 400 time-series. (2) Computing Pearson correlation between each pair of time-series to generate a 400*400 correlation matrix. (3) Pointwise averaging Fisher r-to-z transformed (arctanh) correlation matrices across individuals, and back transforming averages (tanh). In practice, the current computation involved loading in group-average correlation matrices using the BrainSpace function (load_group_fc) and applying the BrainSpace Principal Component Analysis implementation with default parameters. As an aside, the PCA gradients were favored for the current subsequent analyses, over the Diffusion Embedding (Coifman et al. 2005) gradients first derived by Margulies et al. 2016), because they provided a more parsimonious account of semantic variation in the current fMRI data (See **Supplementary Fig. 3**). Nonetheless the two sets of gradients were highly correlated (**Supplementary Fig. 3)**.

To first establish whether any gradient reflected semantic changes in the imagines scenarios, ridge regression was applied to map the participant-specific experiential feature ratings and word embeddings of scenario descriptions to predict gradient responses in new participants and new scenarios. The word-embedding model used was GloVe (Pennington et al. 2014), which represents individual words with three-hundred-dimension feature vectors derived from factoring word co-occurrence counts. These vectors reflect semantics in the sense that words with similar meaning tend to co-occur in natural texts and end up with similar vectors. A “bag-of-words” approach was deployed to model verbal descriptions by a pointwise summation of vectors for constituent words. Although this approach is simple comparative to contemporary Large Language Models because it discounts contextual influences on word meaning, it was favored for the current analyses for interpretability. Specifically, the regression beta-weights on the three hundred features could be interpreted by correlation against a finite dictionary of GloVe embeddings to pinpoint which words drove gradient responses.

### DMN-FPN Gradient#3 activation reflects semantic differences between self-generated mental images

The regression analysis in **Fig. 2** revealed that the DMN-FPN gradient (#3) was the only one predicted with statistical significance, using the experiential and word-embedding models (DMN vs FPN gradient poles are distinguished with blue/red weights). For each gradient and each model, this was evidenced by computing Spearman correlation between predicted and observed data, for ten new scenarios, in a new participant (i.e. both the participant and the scenarios were not present in the training dataset). The generality of effect across the 25 participants was evaluated using signed ranks tests to compare Spearman correlation coefficients to zero (no correlation). The corresponding p-values were corrected for multiple comparisons using False Discovery Rate (Benjamini and Hochberg, 1995). However, although this analysis revealed that DMN-FPN Gradient#3 reflected the semantic content of the imagined scenarios, it did not indicate which semantic features drove prediction.

**Figure 2.**
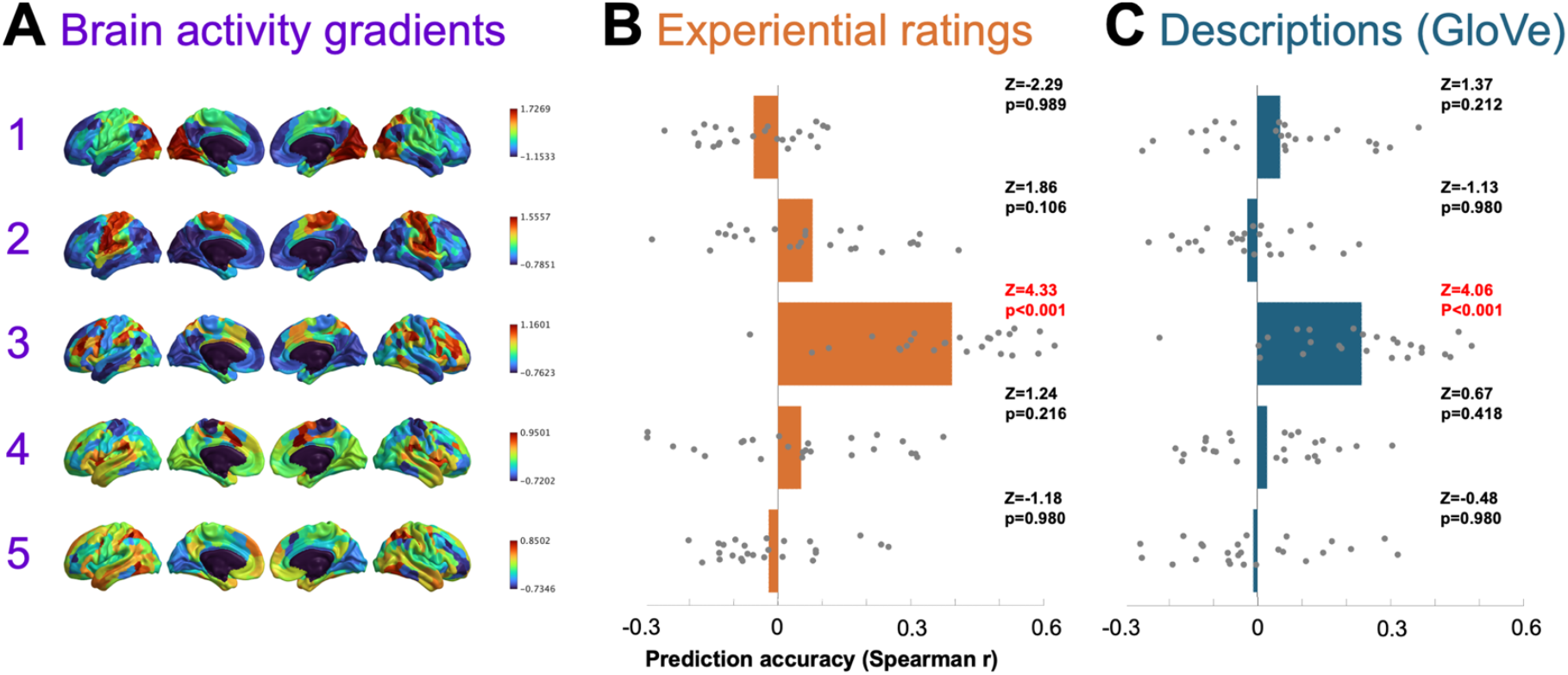
DMN-FPN Gradient#3 projections were selectively predicted by experiential feature ratings and word embedding models of self-reports. **A:** Brain maps of the first five resting-state cortical gradients with weights color-coded from blue (-ve) to red (+ve), see also **Fig. 1A**). **B:** Gradient prediction accuracy using experiential feature ratings. **C:** Gradient prediction accuracy using a GloVe model of participants’ verbal descriptions. In both **B&C** prediction accuracies were estimated using Spearman correlation between predicted and observed gradient activation across the ten held-out test scenarios in held-out participants. Grey dots correspond to prediction accuracies for individual participants and bars are mean prediction accuracies. Signed ranks tests against zero were used to evaluate whether prediction accuracies were significantly greater than zero across the 25 participants. P-values were adjusted according to False Discovery Rate (Benjamini and Hochberg, 1995). Prediction accuracies for Gradient#3 only were estimated to be greater than zero using based on experiential ratings and verbal descriptions. Critically, this suggests a systematic link between Gradient#3 activation and self-generated imagination. Brain images were generated using BrianSpace version 1.10 (Vos de Wael et al. 2020).

### DMN to FPN gradient indexes a transition between imagining social gatherings to solitary activities

To estimate which semantic features underpinned DMN-FPN gradient prediction, we first visualized the profiles of the regression beta-weights on experiential and GloVe features (**Fig. 3**). Because the experiential features each had pre-defined interpretations (**Supplementary Table 1**), the profile of beta-weights were plotted for qualitative inspection (**Fig. 3a**). Features associated with socialization (Social and Communication) had the strongest negative beta-weights – suggesting that they activated the DMN pole of Gradient#3 (which also had negative weights on Gradient#3). Conversely, features associated with action and visuo-tactile perception (Upper Limb, Touch and Bright) were associated with positive beta weights suggesting they activated the Frontoparietal pole which was also positively weighted on Gradient#3 (red).

**Figure 3.**
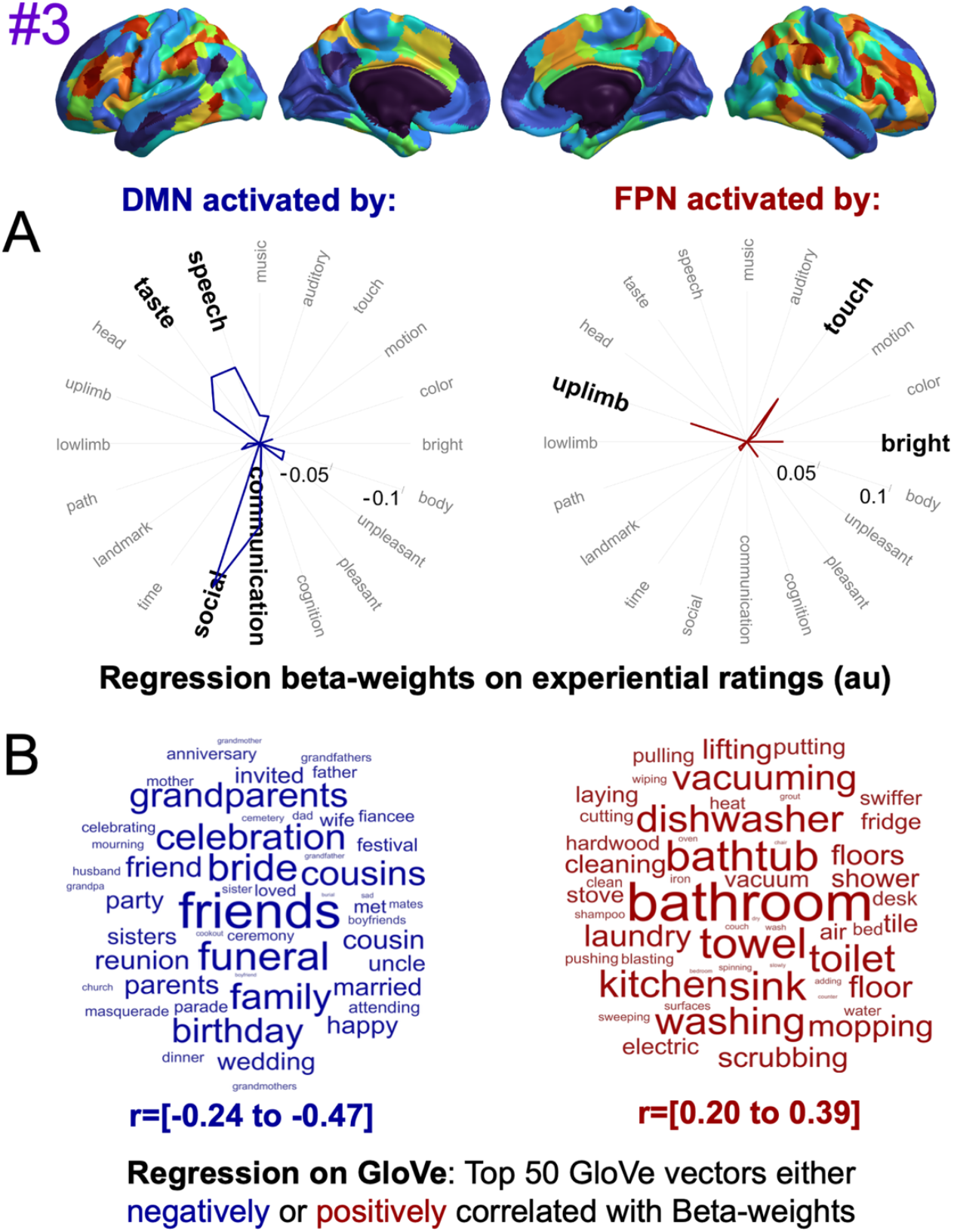
DMN vs FPN activation was predicted by semantic features associated with social vs solitary activities. **A:** Spider plots illustrate Regression Beta-weights assigned to each of the twenty experiential features (averaged across all cross-validated cross-participant regression mappings, see **Fig. 1D**). Axis labels highlighted in bold font correspond to values greater than one SD above the mean. Negative beta-weights (blue) associated with DMN (blue) were strongest for Social and Communication ratings. Positive beta-weights (red) associated with FPN were strongest for Upper-limb and Touch. **B:** Word clouds depicting labels on the top 50 GloVe word vectors that either had the most negative (blue) or most positive (red) correlations with beta-weights on GloVe features. The two analyses suggest that DMN-related blue regions were activated by imagining socializing with friends and family, whereas FPN-related red regions were activated by imagining manual activities conducted alone. Brain images were generated using BrianSpace version 1.10 (Vos de Wael et al. 2020).

Interpretation of the regression beta-weights for the GloVe word embedding vectors - for which features have no prior interpretation - was accomplished by searching for word vectors that were most strongly correlated with the beta-weights on GloVe features. Words with strong positive vs negative correlations would be expected to activate DMN vs FPN respectively. This search was implemented using Spearman correlation to compare the profile of beta-weights across the 300 features to each word in a dictionary formed from the union of 1021 unique words collated across all scenario descriptions provided by the twenty-five participants. **Fig. 3** displays word clouds of the fifty-word vectors with the greatest positive and negative correlations respectively. This analysis suggested that the DMN pole was most strongly activated by words pertaining to friends, family and social events (e.g. celebration, funeral, birthday). In contrast the FPN pole was most strongly activated by words associated with housework, which included rooms (bathroom, kitchen), actions (scrubbing, mopping, vacuuming) and objects (bathtub, dishwasher), but not people. The word embedding results were therefore broadly consistent with the experiential feature ratings in distinguishing socially related features from action and visuo-tactile perception features.

Next, we interpreted DMN-FPN gradient activation, by ranking scenarios according to mean activation values – where scenarios with the smallest negative values, would activate the DMN pole most, and scenarios with the greatest positive values would activate the FPN pole most. Scenario activation was statistically evaluated against a baseline of zero using two-tailed signed rank tests, to detect differences in either negative or positive directions. On the negative end of the scale, and activating the DMN pole, were Party, Funeral, Wedding and Restaurant which all yielded FDR(p)<0.05. At the positive end of the scale, and activating the FPN pole were: Housework, Resting, Exercising, Cooking, Bathing, and Reading, which, with the exception of Bathing [FDR(p)=0.065], all yielded FDR(p)<0.05.

Taken together **Fig. 2** and **3** suggests that the DMN versus FPN were activated when participants imagined social versus solitary activities. However, thus far we have assumed the projection of the imagination data onto Gradient#3 was broadly expressed across multiple subregions of the DMN and FPN, in which case the above claim would hold. However, another possibility was that the effect was only expressed within subregions of one or other network, or in the extreme case, just one single brain region.

### Positive vs negative social correlates were expressed in multiple DMN / FPN subregions

To establish whether the results observed in **Fig. 3/4** broadly generalized across multiple DMN/FPN subregions, and more specifically test the hypothesis that multiple DMN subregions were activated by social relative to solitary scenarios, and multiple FPN regions showed the opposite pattern, we reanalyzed the raw fMRI dataset. To quantify relative activation levels for different scenarios in different brain regions, we estimated hemodynamic response functions (HRF), computed for each scenario in each brain region and each participant.

**Figure 4.**
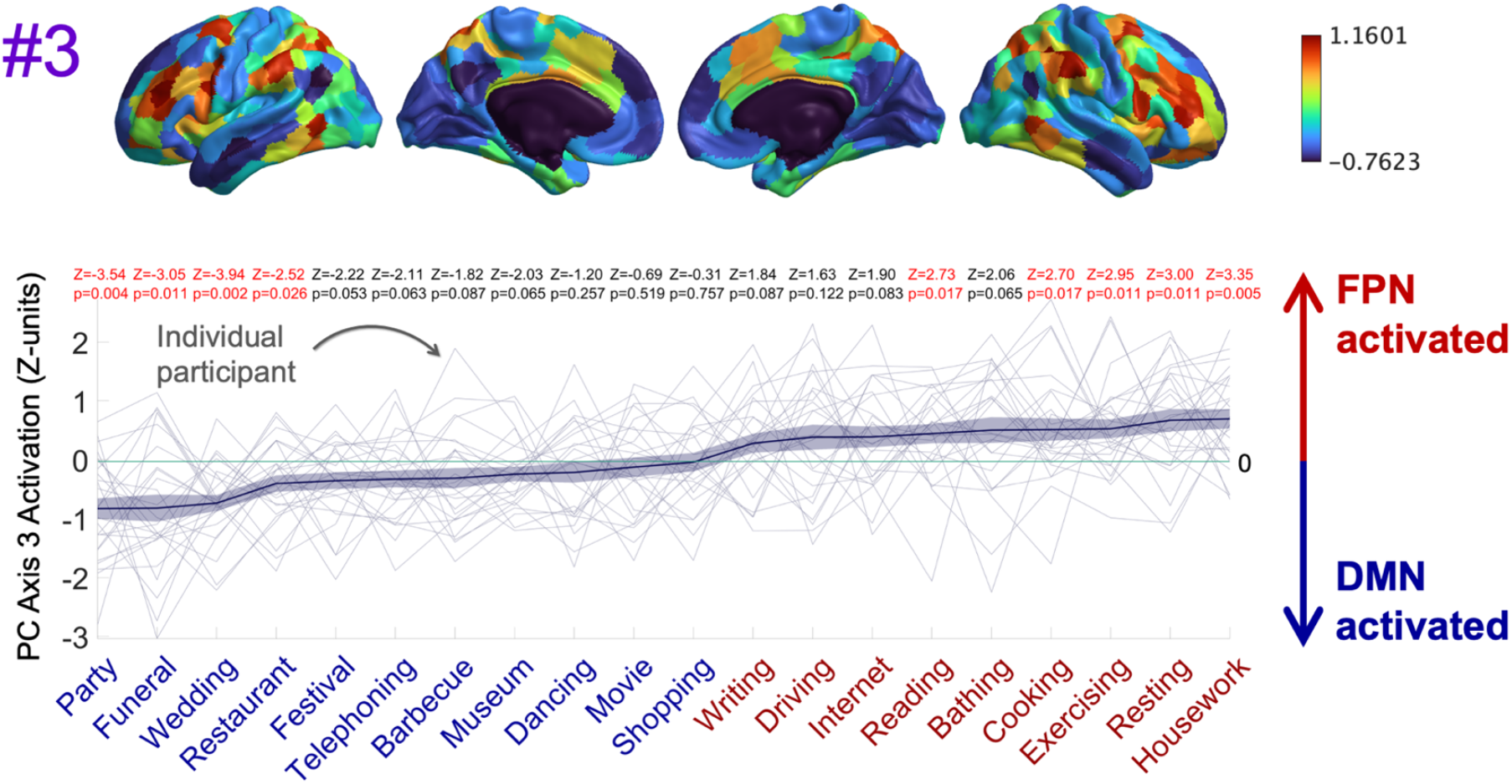
DMN-FPN gradient indexes a transition between imagining more social to more solitary activities. Ranking of scenarios by negative to positive activation on DMN-FPN Gradient #3. **Top:** Cortical map of Gradient#3 with negative-to-positive weight gradient color-coded from blue-to-red. **Bottom:** Scenarios are sorted on the x-axis in ascending order of activation on the Gradient#3 projection. Negative values on Gradient#3 can generally be considered to activate the bluest DMN regions (negative weights) and to deactivate the reddest FPN regions (see also **Fig. 5**). Conversely, positive values on Gradient#3 can generally be considered to activate the reddest FPN regions and deactivate DMN blue ones. The plotted line indicates the Mean+/-SEM Gradient#3 activation across participants. Signed ranks tests against zero were used to evaluate whether individual scenario activation tended to be greater or less than zero across participants (2-tailed). P-values were corrected according to False Discovery Rate (Benjamini and Hochberg, 1995). Party, Funeral, Wedding and Restaurant scenarios yielded values that were significantly below zero (activating the blue DMN-related brain regions). Reading, Cooking, Exercising, Resting and Housework scenarios yielded values that were significantly greater than zero (activating the reddest FPN-related brain regions). The scenario labels on the x-axis are coded blue/red to match the ROIs they activate. The blue/red dichotomy of scenario labels was determined by identifying the zero-crossing point (between Shopping and Writing) and splitting the scenarios accordingly. The same scenario color-codes are applied to color code corresponding hemodynamic response functions (HRFs) in **Fig. 5**. Brain images were generated using BrianSpace version 1.10 (Vos de Wael et al. 2020).

To estimate HRFs, we undertook a series of time-lagged multiple regression analyses to predict fMRI activation in an interval ranging from stimulus presentation time to 15s in the future (the time at which the next stimulus was presented). Thus, each multiple regression had eight predictors [t, t+2.5s, … t+15s] given the TR was 2.5s. Scenario stimulus onset times were indicated by a one in appropriate time-lagged positions in the predictor matrix, and all other values were zero. fMRI and predictor time-series for the entire experiment duration were each z-scored separately such that each time-series had a mean of zero and standard deviation of one. The fitting of each multiple regression produced eight beta-weights, corresponding to each time-lag. For interpretation, a large beta-weight at 7.5s would predict that the corresponding stimulus elicits a strong response with a latency of 7.5s in that specific brain region, scenario and participant. The profile of beta-weights across consecutive time-lags estimates the shape of the hemodynamic response unfolding over time. Illustrative examples of cross-participant HRF profiles estimated for each scenario are displayed in **Fig. 5B**.

**Figure 5.**
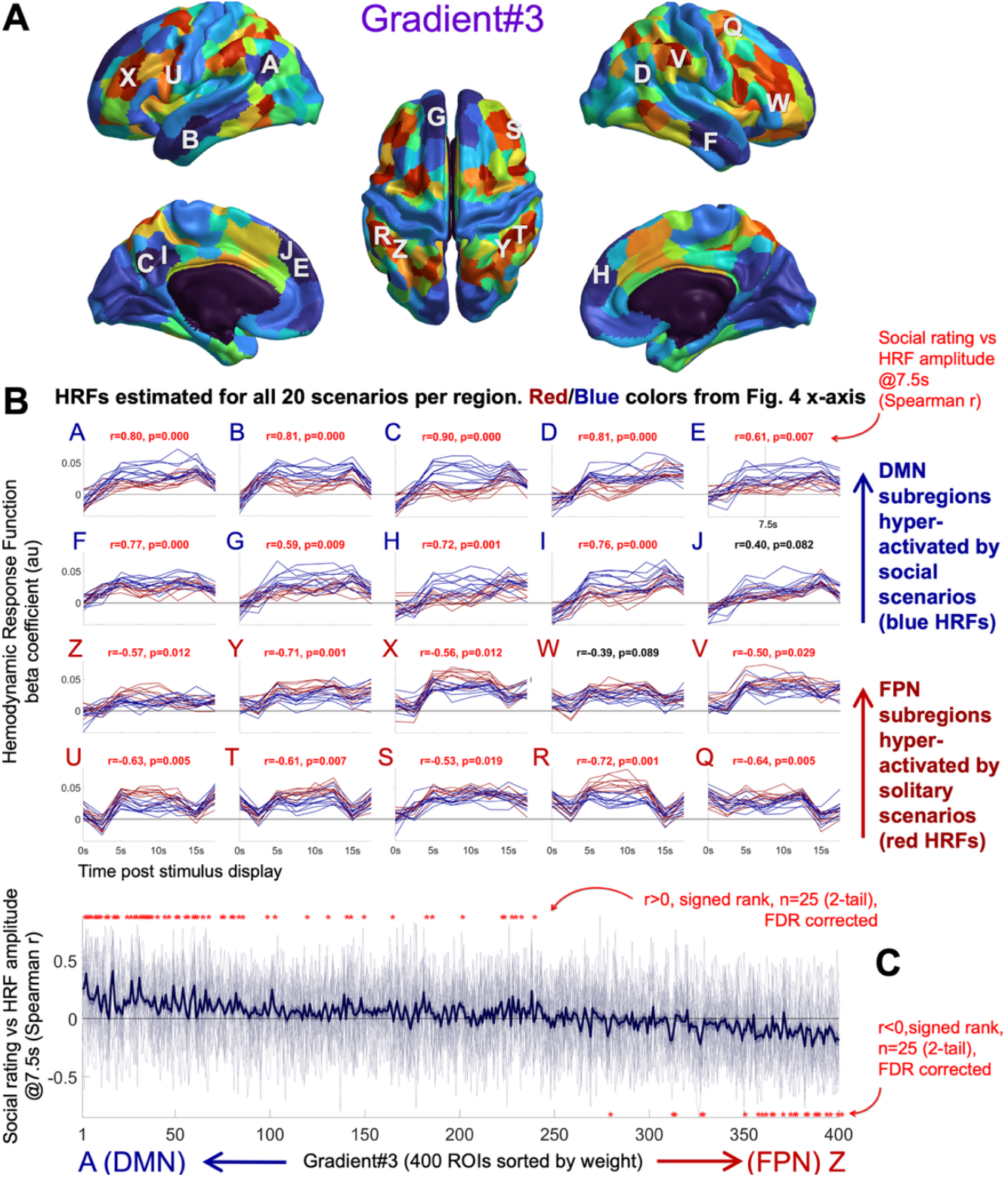
Positive vs negative social correlates were expressed in Multiple DMN vs FPN subregions. **A:** DMN-FPN Gradient#3 with weight gradient color-coded from blue-to-red. Regions A-J (blue) are the ten most negatively weighted regions and Q-Z (red) are the ten greatest positively weighted regions. **B:** Visualization of estimated cross-participant Hemodynamic Response Functions (HRF) in regions A-J, Q-Z, for each of the twenty scenarios. HRFs are colored blue/red according to dichotomy determined on the x-axis of **Fig. 4**. For this visualization, (but not the statistical tests in **C** below) HRFs were computed on data collated across all 25 participants, and HRF amplitudes @7.5s for the twenty scenarios were interpreted with Spearman correlation against the corresponding twenty group average Social ratings. Regions A-J versus Q-Z are broadly distinguished by positive versus negative correlations. **C:** Correlations between participant-specific HRF amplitudes@7.5s and participant-specific Social ratings, estimated for each participant, scenario and brain region (400 total). The thick line is the Mean+/-SEM correlation per region. Brain regions where HRF amplitudes consistently correlated with Social ratings across participants were identified with two-tailed Signed ranks tests against zero (n=25). P-values were adjusted by False Discovery Rate (FDR). Significant positive/negative effects [FDR(p)<0.05] were most concentrated around gradient poles (A,Z) at the left/right of the x-axis. Thus, multiple DMN subregions were hyperactivated by imagining social activities and multiple Frontoparietal subregions were hyper activated by individual activities. Brain images were generated using BrianSpace version 1.10 (Vos de Wael et al. 2020).

To statistically evaluate the hypothesis that HRFs across multiple DMN/FPN subregions would reflect the gradient projection patterns observed in **Fig. 3** and **4**, we analyzed HRF amplitudes at 7.5s. This was on the assumption that amplitudes at 7.5s would be representative of imagination. Here we assumed a standard peak HRF response delay of 4-5s, and on this basis reasoned that 7.5s would give participants >2.5s to both read the stimulus and imagine the corresponding scenario. To interpret HRF amplitudes, we used the corresponding participant-specific Social feature rating as a simple proxy for the social/solitary nature of that scenario, given that social feature ratings (**Fig. 3A**) played a driving role in Gradient#3 prediction (**Fig. 2**). HRF amplitudes were related to Social ratings in the corresponding participants using Spearman correlation. This yielded one correlation coefficient per participant per brain region – where the current would anticipate positive correlation coefficients across DMN subregions (activation levels are greater for social scenarios and weaker for solitary) and negative coefficients in FPN subregions (activation levels are greater for solitary scenarios and weaker for social). To evaluate the generality of outcomes across the 25 participants, two-tailed signed ranks tests were applied to compare the 25 coefficients to zero. Correlation coefficients for each participant across all four hundred brain regions are illustrated in **Fig. 5C** alongside outcomes of the signed rank tests.

The battery of signed rank tests in **Fig. 5C** broadly supported the hypothesis, because significant outcomes [FDR(p)<0.05] tended to cluster around negative (DMN) and positive (FPN) poles of Gradient#3. This provided evidence that: (1) Multiple DMN pole subregions were hyperactivated by social activities. (2) Multiple FPN pole subregions were hyperactivated by solitary activities.

Collectively, these findings show that brain activity elicited in self-generated mental imagery can be characterized along a gradient where opposing activation of DMN versus FPN, differentiates the imagination of social versus individual activities.

### DMN-FPN Gradient#3 activation profiles reflected interparticipant differences in social feature ratings

A final consideration was whether the DMN-FPN Gradient#3 patterns observed in **Fig 2-5** reflected self-generated content as opposed to externally driven semantic content associated with reading the scenario prompts. To test this, we ran an interparticipant differences analysis to evaluate whether participants’ Gradient#3 activation profiles reflected their idiosyncratic Social ratings, reflecting their personal experience over and above other participants. In this analysis, each participant was represented with one vector capturing DMN-FPN Gradient #3 activation values across the twenty scenarios, and another vector capturing their corresponding participant-specific social ratings. Spearman correlation was then computed between gradient and rating vectors, bith within and across participants, yielding a 25 × 25 participant correlation matrix. To assess participant specificity, a rank-sum test compared the 25 congruent within-participant coefficients (on the diagonal) with the 300 unique incongruent coefficients (off-diagonal), under the hypothesis that congruent values would be higher.

This was confirmed (z = 1.89, p = 0.0294), indicating that DMN-FPN activation reflected participant-specific social content, to some degree at least.

To further verify whether interparticipant differences were also reflected in scenarios that drove FPN versus DMN activation, the analysis was repeated on the corresponding nine / eleven scenarios, identified by the red/blue split on the **Fig. 4** x-axis scenario labels. Both tests were significant, suggesting participant-specific content was reflected in each case. Respective signed rank test outcomes were: Z=1.71, p=0.0433 and Z=2.26, p=0.0121.

## Discussion

This study demonstrates that transitions in imagined cognitive states correspond to systematic shifts in brain activity along a default mode–frontoparietal (DMN–FPN) gradient. Frontoparietal regions were more engaged when imagining solitary experiences—ranging from goal-directed activities such as housework and exercising to low action states such as reading—whereas the default mode network was more engaged when imagining social scenarios (e.g., parties). Although this gradient is often described as indexing “task-positive” versus “task-negative” or “external” versus “internal” cognition (Fox et al., 2005), these distinctions do not explain our findings because all states were internally generated – as evidenced by individual differences in personal content, and were elicited by text prompts without explicit goals or interactive demands, beyond reimagining the experience. Instead, our results show that the relative activation levels of both poles of the FPN–DMN gradient track semantic differences in internally generated cognitive states. More broadly, the work establishes a methodological framework to link functional gradients to the semantic content of imagination—which is challenging, since detailed introspective self-reports collected during scanning may disrupt the thoughts they aim to capture.

Conceptually, these findings extend prior work (Spreng et al., 2010; Dixon et al., 2018; Margulies et al., 2016; Huntenburg et al., 2018; Smallwood et al., 2021) in moving beyond the traditional FPN/DMN dichotomy between externally oriented, goal-directed activity (Duncan, 2010; Fedorenko et al., 2013) and internally focused, task-negative processes such as recollection, conceptual integration, and planning (Binder et al., 1999; Mazoyer et al., 2001). They also complement studies showing that activation shifts along the DMN–FPN gradient continuum track on-versus off-task thought and vigilance versus target-detection states (Karapanagiotidis et al., 2020; Turnbull et al., 2020; McKeown et al., 2020). Here, we situate shifts along the same gradient in a richer semantic context, showing that solitary imaginative states recruit frontoparietal regions that overlap those engaged during externally focused, action-oriented tasks, whereas imagined social states resemble off-task or vigilance-related activity, potentially also reflecting social cognition to some degree, at least in off-task thought.

A notable finding was that the frontoparietal system was engaged even in the absence of externally imposed interactive goals typically associated with “task-positive” activation (Duncan, 2010; Fedorenko et al., 2013). As illustrated in **Figs. 4** and **5**, this included imagining both goal-directed solitary activities (e.g., housework, cooking, exercising) and less active states such as reading or resting (which was typically reported as watching TV on the couch, **Supplementary Fig. 2**). Across these conditions, the common feature appeared to be attention to or interaction with non-human objects, suggesting that imagining such interactions may drive frontoparietal engagement, even in the absence of an explicit interactive cognitive task.

Conversely, the default mode network was most strongly engaged during imagined social scenarios such as weddings, funerals, parties, and restaurant visits, consistent with its role in autobiographical memory, social cognition, and theory of mind (Molenberghs et al., 2016; Andrews-Hanna et al., 2010, 2014). These scenarios shared a common emphasis on simulating social environments, which may have included social goals such as a wedding, or internal goals, such as satiating hunger at a restaurant. Together, the findings suggest that the DMN– FPN gradient does not only reflect internal/external or task-positive/task-negative distinctions but also organizes internally generated thought according to whether mental simulations emphasize interaction with social versus non-social environmental factors.

Our framework complements prior research on self-generated thought, which has primarily relied on four methodologies. First, experience-sampling studies ask participants to intermittently report their thoughts (e.g., Delamillieure et al., 2010; Gorgolewski et al., 2014; Smallwood & Schooler, 2015), and fMRI studies have linked DMN activity to off-task (Christoff et al., 2009) and detailed thought (Sormaz et al., 2018), and the DMN-FPN gradient to transitions between external “on-task” states (n-back task events) and unconstrained “off-task” states. However, these designs have limitations: within-scan self-reports are time-consuming, content-focused measures are less reliable than on/off-task judgments (Kane et al. 2021), and the lack of control over off-task states creates substantial interparticipant variability. Second, individual-differences analyses relate resting-state activity to offline trait-level experience sampling reports (Gorgolewski et al., 2014; Smallwood et al., 2016), revealing that many brain regions reflect trait thought patterns, but these correlations do not directly link moment-to-moment changes in brain state to accompanying semantic variations in thought content. Third, reverse-inference approaches infer resting-state network function from separate task-based studies (e.g., Andrews-Hanna et al., 2014; Zhang et al., 2022; Shao et al., 2024), as we do here, but these have lacked detailed measures of self-generated thought content. Fourth, “think-aloud” paradigms link brain activity to real-time verbal reports (Gilmore et al., 2021; Li et al., 2023; Su et al., 2025), but active verbalization can disrupt natural thought, and the timeline of thoughts unfolding may not be accurately reflected by the timing of words spoken. None of these approaches, however, has developed a semantic model capable of predicting gradient responses across both experiential states and individuals, as achieved in the present study.

In conclusion, this work establishes a link between whole-brain activity states in functional gradient space and the imagination of personal experiences. Opposing frontoparietal and default mode network activations correspond to the imagination of solitary versus social scenarios, respectively, showing this gradient does not strictly index external vs internally oriented thought. Our framework provides a means to index how the semantic content of imagination fluctuates moment-to-moment across brain states, and we hope such dynamic measures may in the future complement traditional trait-level resting-state connectivity as a measure of individual differences (Mueller et al., 2013; Seitzman et al., 2016; Tavor et al., 2016).

## Methods

### Participants

The original data collection was approved by the University of Rochester research subject review board (RSRB00067540). All participants received monetary compensation and were required to understand the experimental procedure and give their consent by signing an informed consent form. Data was collected from healthy adults (Mean±SD age=24±3, 16F, 9M).

### Scenario Stimuli

Twenty scenarios were preselected following Anderson et al. (2020) to be diverse events that almost every participant would have personally experienced. Scenarios were presented to participants in the following form: “A X Scenario”, or “An X Scenario” where X is a placeholder for: resting, reading, writing, bathing, cooking, housework, exercising, internet, telephoning, driving, shopping, movie, museum, restaurant, barbecue, party, dancing, wedding, funeral, festival.

### Experimental Procedure

At the beginning of the experiment, an experimenter went through the twenty scenario prompts asking participants to vividly imagine themselves experiencing each scenario, and to actively simulate their sensory experiences, actions and feelings. Once participants had formed a rich mental image, they provided a brief verbal description of their mental image which was transcribed by the experimenter. After all the scenarios had been imagined and described, the experimenter went through the scenarios again, first reminding the participant of their scenario description, and then recording the participant’s ratings of the scenario on 20 experiential features, the likelihood that their mental image reflects a real personal experience (rather than being fictional) and the vividness of the mental image (see **Supplementary Table 1** for the specific instructions for rating each feature, adapted from Binder et al. 2016). The 20 features were a subset of the 65 features collected by Binder et al. 2016 that were chosen to broadly span the twelve domains of experience that were originally identified by Binder et al. 2016 along with the author’s intuitions of which features were most relevant for the 20 scenarios (Anderson et al. 2020). The need to collect 20 rather than all 65 features was mandated by experimental time constraints.

### MRI Data Collection Parameters

Imaging data were collected at the University of Rochester Center for Brain Imaging using a 3T Siemens Prisma scanner (Erlangen, Germany) equipped with a 32-channel receive-only head coil. The fMRI scan began with a MPRAGE scan (TR/TE=1400/2344 ms, TI=702ms, Flip Angle=8°, FOV=256mm, matrix=256×256mm, 192 sagittal slices, slice thickness =1mm, voxel size 1×1×1mm3). fMRI data were collected using a gradient echo-planar imaging (EPI) sequence (TR/TE=2500ms/30ms, Flip Angle=85°, FOV=256mm, 90 axial slices, slice thickness=2mm, voxel size 2×2×2mm3, number of volumes = 639).

### fMRI Experiment

Prior to scanning, participants were reminded of their verbal descriptions of the 20 scenarios (see above) and were requested to vividly reimagine the same mental images in the scanner, when prompted. Outside the scanner they then underwent a single dry run of the fMRI experiment below (a single viewing of all 20 stimuli) to familiarize them with the set up.

In the experiment, stimulus prompts (e.g., “A dancing scenario”) were presented one by one in random order, on a screen in black Arial font (size 50) on a grey background. Prompts remained on screen for 7.5 seconds (3TRs), during which time participants vividly imaged themselves in the scenario. After the prompt was removed there was a 7.5 second delay prior to the next prompt (e.g., “A barbecue scenario”) during which time a fixation cross was displayed. Participants had been instructed to attempt to clear their minds when the prompts were removed.

Participants then underwent a single uninterrupted fMRI session in which the 20 scenario stimuli were presented five times over (five runs). The five repeats later enabled us to compute representations of average mental images (per scenario) and counteract fMRI noise. Stimulus order was randomized within each run. Runs were separated by a 15-second interval, in which a second-by-second countdown was displayed (e.g., “Starting run 2 in 13 seconds”), which was followed by 7.5 seconds of fixation cross preceding the first stimulus of the run. All participants reported that they had been able to imagine the scenarios on prompt.

### MRI Data Preprocessing

MRI data were preprocessed using fMRIPrep (Esteban et al. 2019) to counteract head motion and spatially normalize images to a common neuroanatomical space (MNI152NLin2009cAsym). The boilerplate template detailing the procedure is in **Supplementary Materials**. To counteract nuisance signals and potential confounds in the fMRI data a comprehensive selection of nuisance regressors generated by fMRIPrep were regressed out from each voxels time series. These were 24 head motion parameters (including translation, rotation and their derivatives), white-matter and cerebrospinal fluid timeseries and cosine00, cosine01, cosine02, cosine03. Confound removal was implemented through computing a single multiple regression across the entire fMRI timeline: First, each voxel and each nuisance regressor’s time-series was separately z-scored (by subtracting the mean and dividing by the standard deviation). Second a separate multiple regression was fit for each voxel, mapping nuisance regressors to predict voxel activation. The residuals from the regression (computed separately for each voxel) were taken forward to further analysis.

For analyses, we reduced each participant’s fMRI data to obtain a single volume for each imagined scenario. To do this we first computed a single volume for each scenario replicate, by computing the voxel-wise mean of 4 fMRI volumes (5-15sec) post stimulus onset. To reduce the five replicates of each scenario to a single scenario volume, we again took the voxel-wise mean. This left 20 scenario volumes per individual. The 5 s (2TR) offset was to accommodate hemodynamic response delay, so if a participant were to instantly bring their mental image to mind, the peak response would be around 4-5 seconds (though in practice, mental images are unlikely to have been formed instantly). The four-volume period spans the time until the next stimulus is presented on screen and was set to maximize our chances of capturing the mental image. NB The current averaging approach was selected to accommodate individual differences in the latency and duration of hemodynamic responses associated with internally generated mental images observed by Anderson et al. 2020, which may not be well suited to modeling with a canonical hemodynamic response function time-locked to stimulus presentation.

## Data and Code

Data and code to support analyses are available at DOI: 10.17605/OSF.IO/D4KU8

