## Supplementary Materials for "A default-control network cortical gradient differentiates the imagination of social and solitary experiences"

### Contents:

**Supplementary Figure 1.** Mean experiential feature ratings of imagined scenarios 2

**Supplementary Figure 2.** Word clouds collating content words used to describe imagined scenarios 3

**Supplementary Figure 3**. PCA Gradient #3 was predicted more accurately than Diffusion Embedding gradients and provided a parsimonious account of variation in brain activity that could be predicted by semantic models. 4

**Supplementary Table 1.** Protocol for rating experiential features 5

**fMRIPrep Boilerplate Template** 8

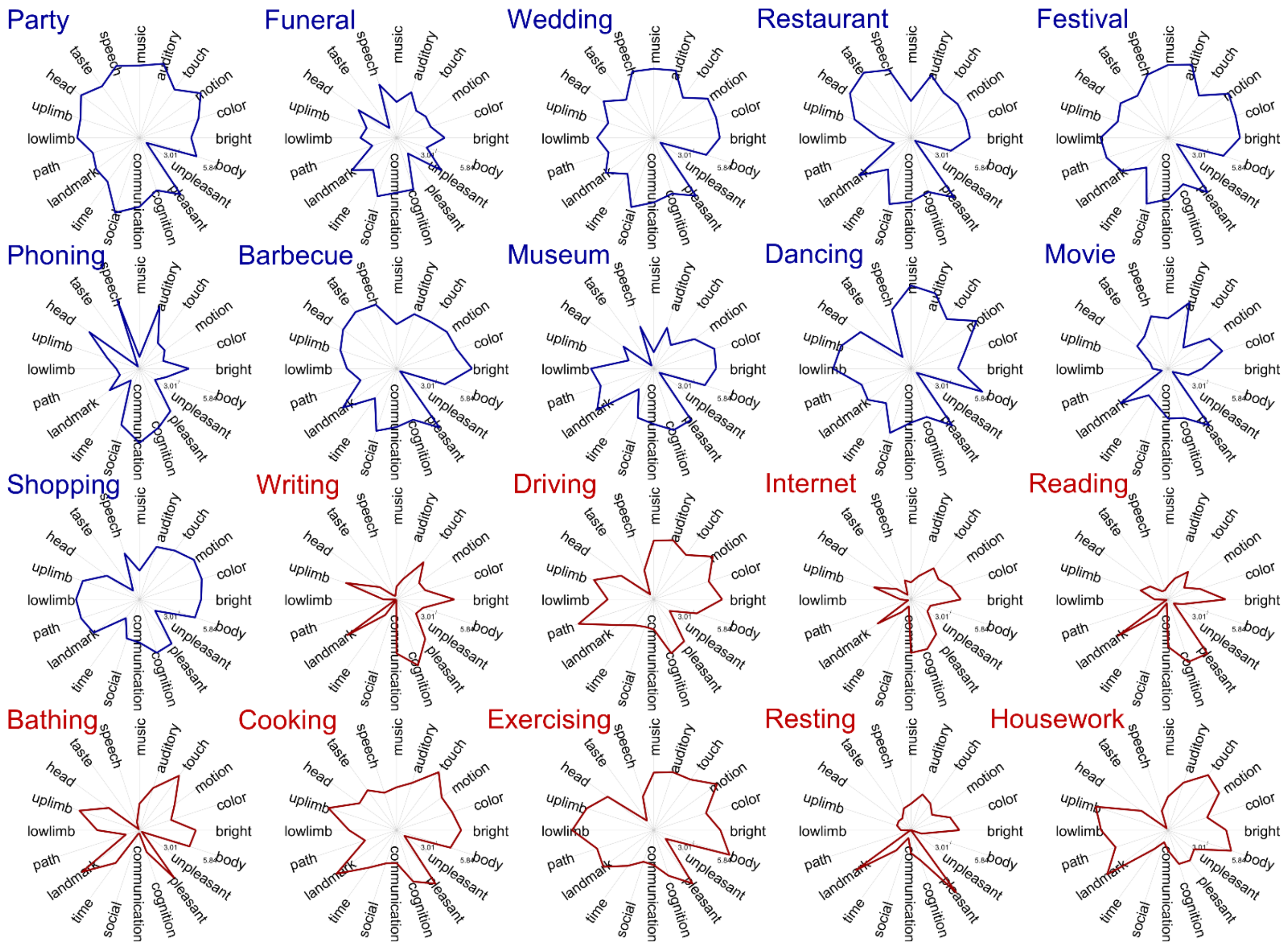


**Supplementary Figure 1**. Mean experiential feature ratings of imagined scenarios

Each spider plot corresponds to participant's self-reports of one scenario (e.g. Party, Housework). Plotted on the twenty axes of each plot are mean participant-specific ratings for each of the twenty experiential features. Red/Blue color coding corresponds to the x-axis scenario labels on **Figure 4**.

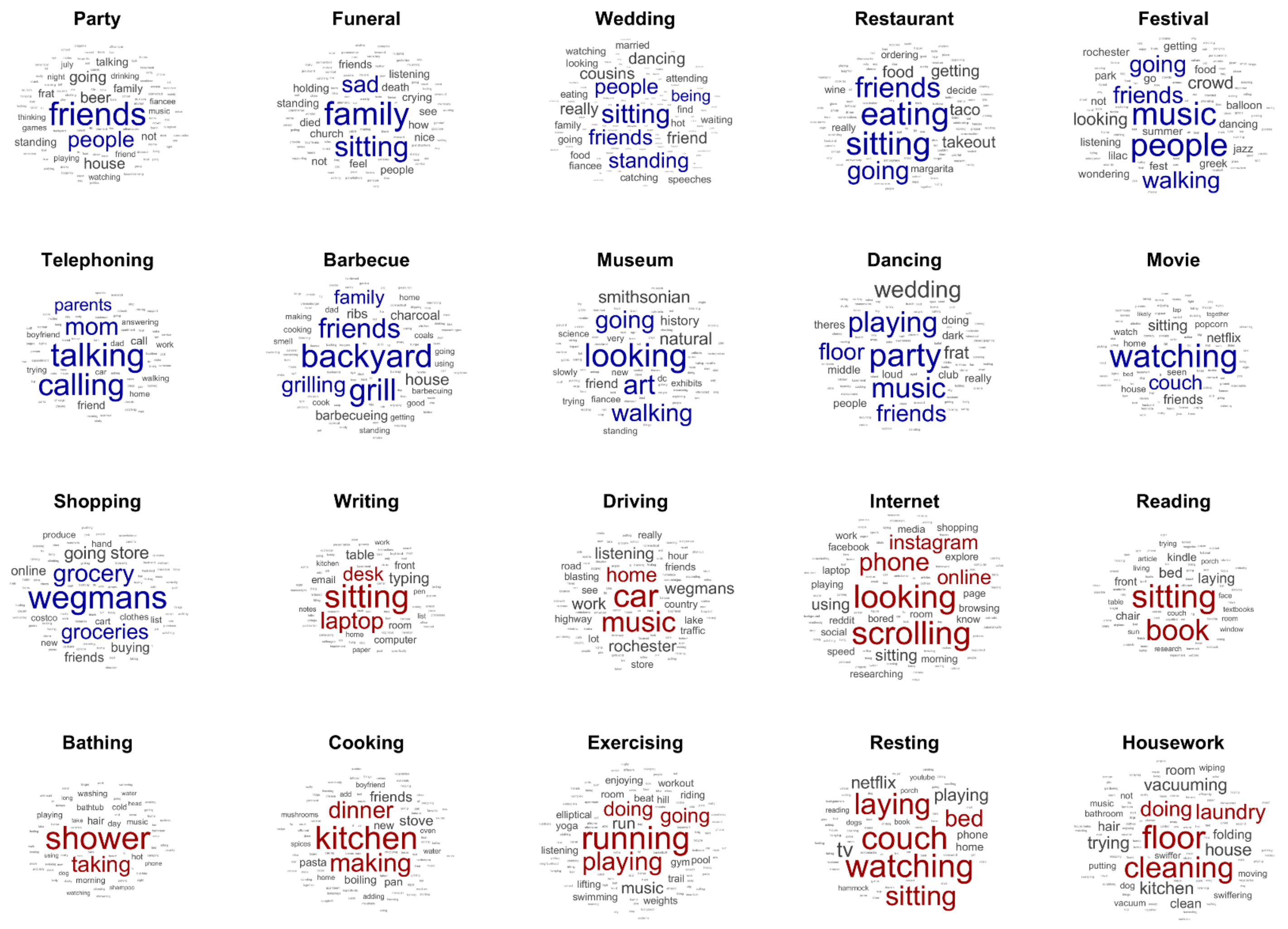


**Supplementary Figure 2**. Word clouds of the content words that were most frequently used to describe each scenario.

Each spider plot corresponds to participant's self-reports of one scenario (e.g., Party, Housework) and displays content words used to describe that scenario, collated across all 25 participants. Font size is scaled by word frequency, and the five most common words, excluding the scenario name itself (e.g. Party) are colored. Word clouds were derived using MATLAB R2023B function wordcloud using default parameters. Red/Blue color coding corresponds to **Figure 4** x-axis scenario labels.

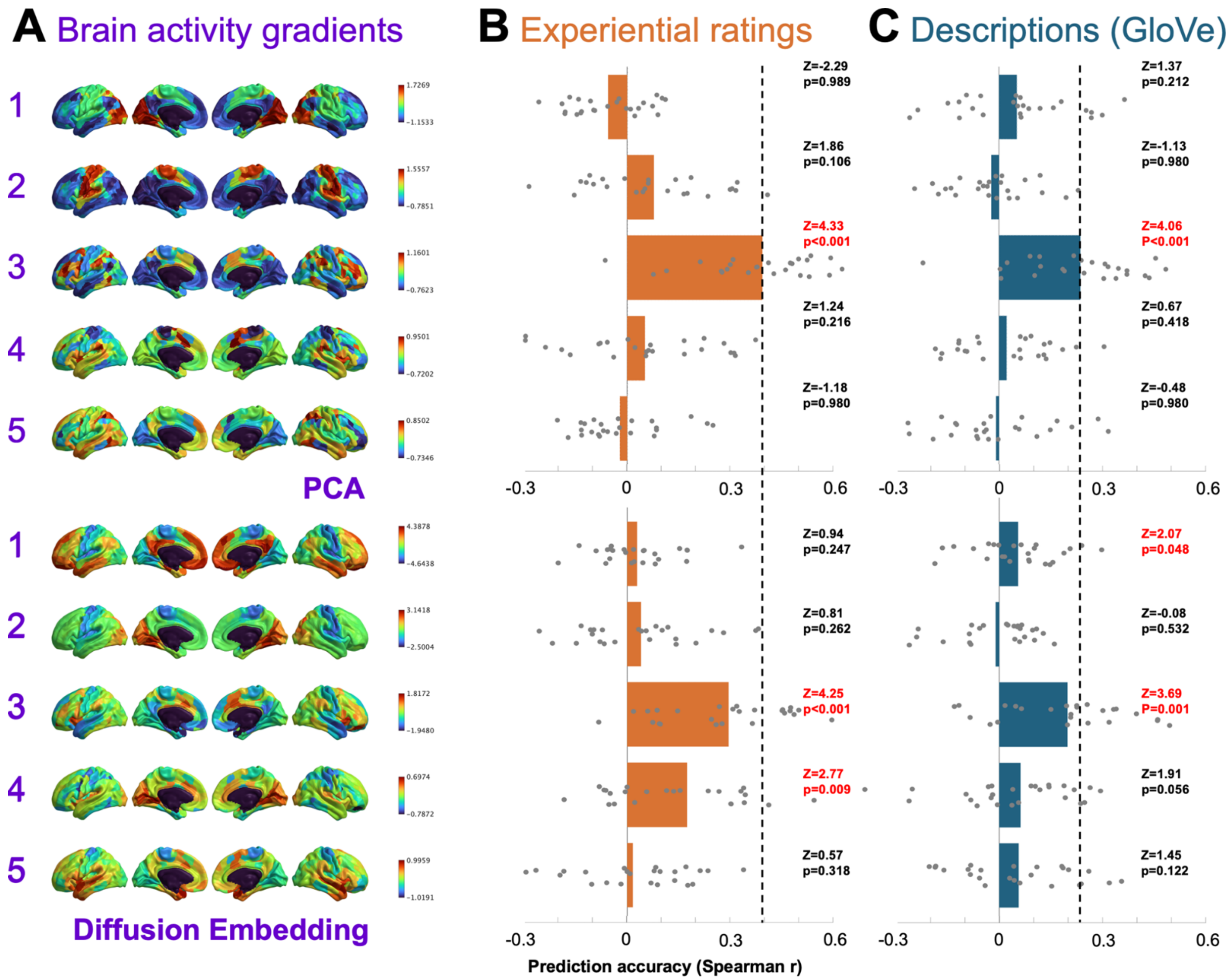


**Supplementary Figure 3**. PCA Gradient #3 more parsimoniously captured semantic variation in fMRI activity than Diffusion Embedding gradients.

PCA gradients[1](Top Row) rather than Diffusion Embedding gradients[2] (Bottom Row) were favored for the current study, because one single PCA gradient (#3) captured semantic variation in the current fMRI dataset, and this gradient was predicted with relatively high accuracy comparative to all other gradients tested.

The PCA analysis in the top row is precisely the same as **Fig. 3**. The same analysis is recomputed using projections on Diffusion Embedding gradients[2] from Neurovault[3] (bottom row). Each gradient was represented as voxel averages within Schaefer-400[4] regions. Brain images were generated using BrainSpace v 1.10[5].

**A**. Gradients derived by PCA and Diffusion Embeddings. **B:** Gradient prediction accuracy using experiential feature ratings. **C:** Gradient prediction accuracy using a GloVe model of participants' verbal descriptions. In **B&C**, prediction accuracies were estimated using Spearman correlation between predicted and observed gradient activation across the ten held-out test scenarios in held-out participants. Grey dots correspond to prediction accuracies for individual participants and bars are mean prediction accuracies. Signed ranks tests against zero were used to evaluate whether prediction accuracies were significantly greater than zero across the 25 participants. P-values were adjusted according to False Discovery Rate (FDR[6]).

PCA and Diffusion embedding gradients #1 to #5 were correlated at Pearson r=[-0.65, -0.78, 0.80, 0.51, 0.32].

**Supplementary Table 1.** Protocol for rating 20 experiential features (black font), and vividness and likelihood (blue font).

| | |
|---|---|
| **Bright** | Please rate the degree to which your imagined scenario contains light or brightness. Example high score: 6; The sun is very bright. Example medium score: 3; When something glows it gives off a soft light. Example low score: 0; When something gives off no light. |
| **Color** | Please rate to what degree you think of each scenario as involving color or changes in color. Example high score: 6; To blush is to turn red. Example medium score: 3; Decorating can involve adding or changing a color. Example low score: 0; Color does not play a role. |
| **Motion** | Please rate to what degree you think of each scenario as involving a specific type or a large amount of visible movement. Example high score: 6; When something bounces it is constantly moving. Example medium score: 3; Clock parts move, but not very noticeably. Example low score: 0; The scene is still. |
| **Touch** | Please rate to what degree each of your scenarios involves an action or activity in which something is felt by touch. Example high score: 6; Caressing means touching something softly. Example medium score: 3; Perceiving something may in some cases involve touching it. Example low score: 0; Touch does not play a role. |
| **Audition** | Please rate to what degree each of your scenarios involves hearing something. Example high score: 6; When something beeps it makes a sound. Example medium score: 3; Perceiving something may involve sound. Example low score: 0; The scenario involves no sound. |
| **Music** | Please rate to what degree each of your scenarios involves music or musical sounds. Example high score: 6; Singing creates musical sounds. Example medium score: 3; Chiming noises can be somewhat musical. Example low score: 0; The scenario involves no musical sounds |
| **Speech** | Please rate the degree to which each of your scenarios involves human speech sounds. Example high score: 6; Talking involves human speech sounds. Example medium score: 3; Babbling often refers to human speech that is hard to understand. Example low score: 0; The scenario involves no human speech sounds. |
| **Taste** | Please rate to what degree each of your scenarios involves tasting something. Example high score: 6; Sipping involves tasting a beverage. Example medium score: 3; Cooking is often accompanied by tasting. Example low score: 0; Taste does not play a role in this scenario. |

| | |
|---|---|
| **Head** | Please rate to what degree each of your scenarios involves the use of the face, mouth, or tongue. Example high score: 6; Smiling is an action involving the face and mouth. Example medium score: 3; Breathing involves the mouth or nose, although they don't actually move. Example low score: 0; The head does not play a role in this scenario. |
| **Upper Limbs** | Please rate to what degree each of your scenarios involves the use of the arms, hands, or fingers. Example high score: 6; Applauding is an action involving the arms and hands. Example medium score: 3; Jogging usually involves the arms to some degree. Example low score: 0; The upper limbs do not play a role in this scenario. |
| **Lower Limbs** | Please rate to what degree each of your scenarios involves the use of the leg(s) or feet. Example high score: 6; Jumping requires using your legs and feet. Example medium score: 3; Sitting involves some minimal positioning of the legs. Example low score: 0; The lower limbs do not play a role in this scenario. |
| **Path** | Please rate to what degree each of your scenarios involves someone or something moving from one location to another. Example high score: 6; Traveling involves going from one place to another. Example medium score: 3; Searching may or may not require you to change your location. Example low score: 0; The scenario does not involve moving around. |
| **Landmark** | Please rate to what degree each of your scenarios involves an action or activity that occurs at a fixed location, as on a map. Example high score: 6; Libraries and other buildings have a very fixed location. Example medium score: 3; Bushes have a fixed location but are not distinctive enough to be marked on maps. Example low score: 0; The imagined scenario could happen anywhere. |
| **Time** | Please rate to what degree each of your scenarios involves an occurrence at a typical or predictable time. Example high score: 6; Waking up is something you do at a certain time of the day. Example medium score: 3; Cooking is something that often occurs in the evening. Example low score: 0; The scenario does not occur at a specific time. |
| **Social** | Please rate to what degree each of your scenarios involves interactions between people. Example high score: 6; Collaborating requires interactions between people. Example medium score: 3; Driving in a car is often done with other people. Example low score: 0; The scenario does not involve other people. |
| **Communication** | Please rate to what degree each of your scenarios involves communication or transmitting/receiving information. Example high score: 6; Explaining is when a person clarifies by communicating information. Example medium score: 3; Painting may be an |

| | |
|---|---|
| | artistic form of communication. Example low score: 0; The scenario does not involve communication. |
| **Cognition** | Please rate to what degree each of your scenarios involve a mental activity or state of mind that involves thinking. Example high score: 6; Considering something involves thinking about it. Example medium score: 3; Grieving is a state of mind that involves some degree of thinking. Example low score: 0; The scenario does not involve thinking. |
| **Pleasant** | Please rate to what degree each of your scenarios involves something that is pleasant. Example high score: 6; Relaxing is probably something you find pleasant. Example medium score: 3; Conversing is probably something you find somewhat pleasant. Example low score: 0; The scenario does not involve anything pleasant. |
| **Unpleasant** | Please rate to what degree each of your scenarios involves something that is unpleasant. Example high score: 6; Arguing is something you probably find unpleasant. Example medium score: 3; waiting is probably something you find somewhat unpleasant. Example low score: 0; the scenario does not involve anything unpleasant. |
| **Body** | Please rate to what degree each of your scenarios involves visible movements of the body or limbs. Example high score: 6; Running produces visible movements of the body. Example medium score: 3; Shuddering produces small visible movements of the body. |
| **Vividness** | Please rate how vivid, or how well you are able to picture each of your scenarios. Example high score: 6; Very vivid mental image. Example medium score: 3; Some mental image. Example low score: 0; Not able to create a vivid mental image. |
| **Likelihood** | Please rate the likelihood of each scenario having happened, or how likely it is to happen. Example high score: 6; This scenario has happened to me. Example medium score: 3; It might or could happen. Example low score: 0; This is a fictitious situation. |

**fMRIPrep Boilerplate Template**

### Copyright Waiver

The below boilerplate text was automatically generated by fMRIPrep with the express intention that users should copy and paste this text into their manuscripts unchanged. It is released under the CC0 license.

### Boilerplate

Results included in this manuscript come from preprocessing performed using fMRIPrep 20.2.1 (Esteban, Markiewicz, et al. (2018)[7]; Esteban, Blair, et al. (2018)[8]; RRID:SCR_016216), which is based on Nipype 1.5.1 (Gorgolewski et al. (2011)[9]; Gorgolewski et al. (2018)[10]; RRID:SCR_002502).

### Anatomical data preprocessing

A total of 1 T1-weighted (T1w) images were found within the input BIDS dataset.The T1-weighted (T1w) image was corrected for intensity non-uniformity (INU) with N4BiasFieldCorrection (Tustison et al. 2010[11]), distributed with ANTs 2.3.3 (Avants et al. 2008[12], RRID:SCR_004757), and used as T1w-reference throughout the workflow. The T1w-reference was then skull-stripped with a Nipype implementation of the antsBrainExtraction.sh workflow (from ANTs), using OASIS30ANTs as target template. Brain tissue segmentation of cerebrospinal fluid (CSF), white-matter (WM) and gray-matter (GM) was performed on the brain-extracted T1w using fast (FSL 5.0.9, RRID:SCR_002823, Zhang, Brady, and Smith 2001[13]). Brain surfaces were reconstructed using recon-all (FreeSurfer 6.0.1, RRID:SCR_001847, Dale, Fischl, and Sereno 1999[14]), and the brain mask estimated previously was refined with a custom variation of the method to reconcile ANTs-derived and FreeSurfer-derived segmentations of the cortical gray-matter of Mindboggle (RRID:SCR_002438, Klein et al. 2017[15]). Volume-based spatial normalization to one standard space (MNI152NLin2009cAsym) was performed through nonlinear registration with antsRegistration (ANTs 2.3.3), using brain-extracted versions of both T1w reference and the T1w template. The following template was selected for spatial normalization: ICBM 152 Nonlinear Asymmetrical template version 2009c [Fonov et al. (2009)[16], RRID:SCR_008796; TemplateFlow ID: MNI152NLin2009cAsym],

### Functional data preprocessing

For each of the 1 BOLD runs found per subject (across all tasks and sessions), the following preprocessing was performed. First, a reference volume and its skull-stripped version were generated using a custom methodology of fMRIPrep. Susceptibility distortion correction (SDC) was omitted. The BOLD reference was then co-registered to the T1w reference using bbregister (FreeSurfer) which implements boundary-based registration (Greve and

Fischl 2009[17]). Co-registration was configured with six degrees of freedom. Head-motion parameters with respect to the BOLD reference (transformation matrices, and six corresponding rotation and translation parameters) are estimated before any spatiotemporal filtering using mcflirt (FSL 5.0.9, Jenkinson et al. 2002[18]). BOLD runs were slice-time corrected using 3dTshift from AFNI 20160207 (Cox and Hyde 1997[19], RRID:SCR_005927). The BOLD time-series (including slice-timing correction when applied) were resampled onto their original, native space by applying the transforms to correct for head-motion. These resampled BOLD time-series will be referred to as preprocessed BOLD in original space, or just preprocessed BOLD. The BOLD time-series were resampled into standard space, generating a preprocessed BOLD run in MNI152NLin2009cAsym space. First, a reference volume and its skull-stripped version were generated using a custom methodology of fMRIPrep. Several confounding time-series were calculated based on the preprocessed BOLD: framewise displacement (FD), DVARS and three region-wise global signals. FD was computed using two formulations following Power (absolute sum of relative motions, Power et al. (2014)[20]) and Jenkinson (relative root mean square displacement between affines, Jenkinson et al. (2002)[18]). FD and DVARS are calculated for each functional run, both using their implementations in Nipype (following the definitions by Power et al. 2014[20]). The three global signals are extracted within the CSF, the WM, and the whole-brain masks. Additionally, a set of physiological regressors were extracted to allow for component-based noise correction (CompCor, Behzadi et al. 2007[21]). Principal components are estimated after high-pass filtering the preprocessed BOLD time-series (using a discrete cosine filter with 128s cut-off) for the two CompCor variants: temporal (tCompCor) and anatomical (aCompCor). tCompCor components are then calculated from the top 2% variable voxels within the brain mask. For aCompCor, three probabilistic masks (CSF, WM and combined CSF+WM) are generated in anatomical space. The implementation differs from that of Behzadi et al. in that instead of eroding the masks by 2 pixels on BOLD space, the aCompCor masks are subtracted a mask of pixels that likely contain a volume fraction of GM. This mask is obtained by dilating a GM mask extracted from the FreeSurfer's aseg segmentation, and it ensures components are not extracted from voxels containing a minimal fraction of GM. Finally, these masks are resampled into BOLD space and binarized by thresholding at 0.99 (as in the original implementation). Components are also calculated separately within the WM and CSF masks. For each CompCor decomposition, the k components with the largest singular values are retained, such that the retained components' time series are sufficient to explain 50 percent of variance across the nuisance mask (CSF, WM, combined, or temporal). The remaining components are dropped from consideration. The head-motion estimates calculated in the correction step were also placed within the corresponding confounds file. The confound time series derived from head motion estimates and global signals were expanded with the inclusion of temporal derivatives and quadratic terms for each (Satterthwaite et al. 2013[22]). Frames that exceeded a threshold of 0.5 mm FD or 1.5 standardised DVARS were annotated as motion outliers. All resamplings can be performed with a single interpolation step by composing all the pertinent transformations (i.e. head-motion transform matrices, susceptibility distortion correction when available, and co-registrations to anatomical and output spaces). Gridded (volumetric) resamplings were performed using antsApplyTransforms (ANTs), configured with Lanczos interpolation to minimize the smoothing effects of other kernels (Lanczos 1964[23]). Non-gridded (surface) resamplings were performed using mri_vol2surf (FreeSurfer).

Many internal operations of fMRIPrep use Nilearn 0.6.2 (Abraham et al. 2014[24], RRID:SCR_001362), mostly within the functional processing workflow. For more details of the pipeline, see the section corresponding to workflows in fMRIPrep's documentation.
